# Proteomic Screening Identifies HNRNPA2B1 as an Epigenetic Repressor of Epstein-Barr Virus Reactivation

**DOI:** 10.64898/2026.02.27.708571

**Authors:** Febri Gunawan Sugiokto, Yuxin Liu, Renfeng Li

**Author notes:** Corresponding author (RL).

## Abstract

Epstein-Barr virus (EBV) establishes lifelong persistent infection in over 90% of the world’s population. The virus persists as an episome in the host cells during latency and periodically reactivates through transcriptional activation of the immediate-early (IE) genes. While epigenetic regulation is central to maintaining viral latency, the host factors that enforce repression at these promoters remain incompletely defined. Here, we employed a novel Clustered Regularly Interspaced Short Palindromic Repeats (CRISPR)/dCas9-based <u>en</u>gineered DNA-binding molecule-mediated <u>ch</u>romatin immuno<u>p</u>recipitation coupled with <u>m</u>ass <u>s</u>pectrometry (enChIP-MS) approach to identify proteins associated with the promoter of EBV IE gene *ZTA*. This approach revealed an enrichment of multiple heterogeneous nuclear ribonucleoproteins and identified HNRNPA2B1 as a potential regulator of EBV *ZTA* gene expression. Functional analyses across multiple EBV+ cancer cell models demonstrated that HNRNPA2B1 acts as a restriction factor for EBV lytic reactivation. Depletion of HNRNPA2B1 led to increased expression of IE and downstream lytic genes, enhanced RNA polymerase II recruitment to the *ZTA* and *RTA* promoters, and elevated the proportion of cells entering the lytic cycle. Conversely, enforced expression of HNRNPA2B1 suppressed EBV lytic reactivation. Mechanistically, HNRNPA2B1 enhances repressive viral chromatin states by facilitating recruitment of the histone demethylase KDM1A to EBV IE gene promoters, thereby limiting the activating histone H3 lysine 4 trimethylation. Together, these findings identify HNRNPA2B1 as a key epigenetic regulator of EBV latency and link RNA-binding proteins to epigenetic control of viral reactivation.

**IMPORTANCE:** This study identifies HNRNPA2B1 as a previously unrecognized host factor that promotes Epstein-Barr virus (EBV) latency through direct regulation of viral chromatin at immediate-early gene promoters. By integrating locus-specific chromatin proteomics with functional and mechanistic analyses, our work reveals how an RNA-binding protein HNRNPA2B1 recruits a histone-modifying enzyme to control EBV reactivation. These findings provide new insights into host-virus interactions that controls EBV latency and reactivation and highlight the role of RNA-binding proteins in chromatin regulation that may be broadly relevant to other latent DNA viruses.

## INTRODUCTION

Epstein-Barr virus (EBV) is a ubiquitous human gamma-herpesvirus that establishes lifelong latency in infected cells [1-4]. During latency, the viral genome persists as a chromatinized episome that is subject to host epigenetic regulation [5-7]. Tight repression of viral lytic gene expression is essential for immune evasion and long-term persistence, whereas disruption of this control promotes lytic reactivation and contributes to viral dissemination and disease [1]. Central to this process is transcriptional regulation of the EBV immediate-early (IE) genes *ZTA* and *RTA*, which function as master regulators that initiate the lytic transcriptional cascade required for viral DNA replication, capsid assembly, and the release of progenies [8-12].

The repressive chromatin state of the *ZTA* and *RTA* promoters, namely ZTAp and RTAp, plays a critical role in controlling EBV latency. In latently infected cells, these promoters are maintained in a transcriptionally restrictive state characterized by low RNA polymerase II occupancy and limited enrichment of activating histone modifications [13, 14]. Among these, trimethylation of histone H3 lysine 4 (H3K4Me3) is a key determinant of promoter activation [15]. H3K4Me3 is broadly associated with transcriptionally active genes and facilitates recruitment of the transcriptional machinery. During EBV lytic reactivation, rapid accumulation of H3K4Me3 at the *ZTA* and *RTA* promoters accompanies transcriptional activation, whereas removal of this mark promotes latency [14].

Dynamic regulation of H3K4 methylation is controlled by opposing activities of histone methyltransferases and lysine demethylases [16]. A recent study showed that the histone demethylase KDM1A (LSD1) acts as a critical restriction factor for EBV lytic replication by removing activating H3K4Me3 mark to maintain repressive viral chromatin [17].

RNA binding proteins have increasingly been recognized as important regulators of viral infection beyond their canonical roles in RNA processing. Heterogeneous nuclear ribonucleoproteins are multifunctional factors that coordinate transcription, RNA metabolism, and chromatin organization. Among them, HNRNPA2B1 has been implicated in the life cycles of multiple viruses, where it can influence viral RNA stability, nuclear export or host immune response to viral infection [18-26]. HNRNPA2B1 has also been linked to transcriptional regulation in host cells, where it acts at the interface of RNA regulation and epigenetic control of chromatin through lncRNA, *HOTAIR* [27, 28]. Despite these emerging roles, whether HNRNPA2B1 contributes directly to the repression of viral chromatin in latently infected cells has not been defined.

Recent advances in Clustered Regularly Interspaced Short Palindromic Repeats (CRISPR)-based chromatin targeting have enabled locus-specific identification of proteins associated with defined genomic regions. This approach employs catalytically inactive Cas9 (dCas9) fused to an epitope tag, such as Flag or HA, in a strategy termed <u>en</u>gineered DNA-binding molecule-mediated <u>ch</u>romatin immuno<u>p</u>recipitation (enChIP). Targeting dCas9 to a specific genomic locus allows affinity purification of the associated chromatin and interacting proteins, which can subsequently be identified by mass spectrometry (MS) [29-32]. Applying enChIP-MS to viral episome provides a powerful strategy to uncover host factors that directly regulate viral chromatin. From our enChIP-MS analysis, we discovered that a group of heterogeneous nuclear ribonucleoproteins (HNRNPs) were significantly enriched at EBV ZTAp. In this study we focused on HNRNPA2B1 for further analysis of its role in EBV latency and reactivation. Our study revealed that HNRNPA2B1 represses EBV IE promoters by recruitment of the histone demethylase KDM1A, thus limiting H3K4Me3 accumulation, promoting latency and restricting lytic reactivation. These results establish a previously unrecognized role for HNRNPA2B1 in viral epigenetic regulation and highlight a mechanism linking RNA binding proteins to chromatin-based control of EBV latency and reactivation.

## RESULTS

### Identification of HNRNPs as EBV promoter binding proteins using enChIP-MS

Previously, we developed a CRISPR-mediated EBV reactivation (CMER) system using a dCas9-based strategy in which dCas9 was fused to the transcriptional activator VP64 and directed to EBV ZTAp to induce EBV lytic reactivation [33]. To extend the utility of this platform, we cloned a non-targeting control sgRNA and a ZTAp-targeting sgRNA (sg5) into the CRISPR/dCas9-Flag vector for enChIP analysis of ZTAp using Akata (EBV+) Burkitt lymphoma cells.

By employing enChIP-MS, we sought to identify proteins enriched at the ZTAp that may function as transcriptional repressors. In this system, Flag-tagged dCas9 is directed to the ZTAp of the EBV episome, enabling selective enrichment and identification of chromatin-associated proteins at this regulatory region. Following crosslinking, chromatin fragmentation, and Flag-based ChIP, the ZTAp-associated proteins will be subjected to trypsin digestion and then analyzed by Liquid Chromatography-Tandem Mass Spectrometry (LC-MS/MS) (**Fig. 1A**).

**Figure 1.**
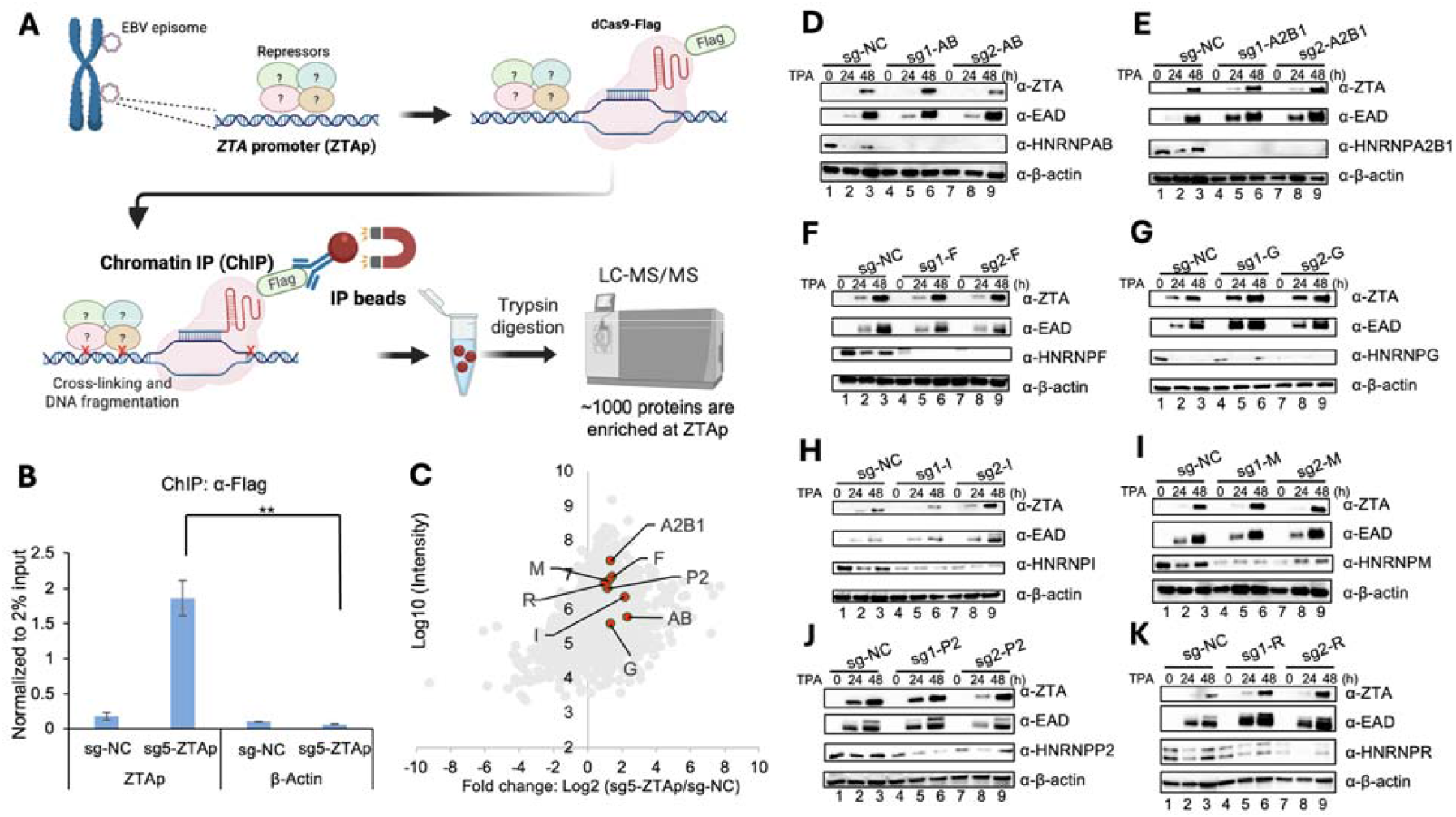
Identification of HNRNP proteins as repressors of EBV lytic reactivation using enChIP-MS and targeted CRISPR/Cas9-based gene knockout screening. (**A**) Schematic illustration of the experimental workflow of enChIP-MS. Flag-tagged dCas9 is guided to the ZTAp on the EBV episome using a ZTAp-specific sgRNA. Chromatin is crosslinked and fragmented, followed by Flag-based ChIP as described in Material and Methods. ChIPed proteins are digested with trypsin and analyzed by LC-MS/MS to identify proteins enriched at the ZTAp. (**B**) Akata (EBV+) B cells were used to create cell lines using lentivirus carrying dCas9-Flag with control (sg-NC) and ZTAp-targeting sgRNA (sg5-ZTAp). The enrichment of Flag-dCas9 at the ZTAp and a control β-actin locus was quantified by ChIP-qPCR and normalized to 2% input as described in Material and Methods. Data are shown as mean ± SD. **p<0.01. (**C**) Scatter plot showing fold change (log_2_ sg5-ZTAp/sg-NC) versu signal intensity (log_10_) of proteins identified by enChIP-MS. Eight HNRNPs with greater than 2-fold enrichment at the ZTAp were highlighted. (**D-K**) WB analysis of EBV lytic proteins (ZTA and EAD) and HNRNPs in EBV+ SNU-719 cells following depletion of individual HNRNP family members and then lytically induced by TPA treatment for 0, 24 and 48 h. β-actin serves as a loading control.

ChIP followed by quantitative polymerase chain reaction (ChIP-qPCR) analysis confirmed efficient and specific targeting of dCas9 to the ZTAp, with significantly greater enrichment observed in cells expressing the ZTAp-targeting sgRNA (sg5-ZTAp) compared with the non-targeting control sgRNA (sg-NC), while no enrichment was detected at the β*-actin* locus (**Fig. 1B**). Proteomic analysis identified approximately 1,000 proteins enriched at the ZTAp using twofold enrichment cut-off relative to sg-NC. Interestingly, eight HNRNPs were significantly enriched at the ZTAp, suggesting a potential role for RNA-binding proteins in regulating EBV lytic gene expression (**Fig. 1C**).

To functionally evaluate the role of these HNRNPs in EBV lytic reactivation, individual HNRNP was depleted using CRISPR/Cas9-mediated gene targeting in EBV+ gastric carcinoma cells SNU-719 (**Fig. 1D-K**) [34], an epithelial cell line amenable for CRSPR/Cas9 screening of multiple candidate genes with higher efficiency than B cells. Western blot (WB) analysis revealed that depletion of HNRNPA2B1 (**Fig. 1E**), HNRNPG (**Fig. 1G**), and HNRNPR (**Fig. 1K**) resulted in significant increased expression of ZTA and the early lytic protein EAD compared to the non-targeting control cells following lytic induction by 12-O-tetradecanoylphorbol-13-acetate (TPA) treatment, whereas depletion of other HNRNPs (AB/F/I/M/P2) did not affect lytic induction (**Fig. 1D, 1F, 1H, 1I and 1J**). These results suggest that multiple HNRNP proteins function as restriction factors that limit EBV lytic gene expression.

### HNRNPA2B1 functions as a conserved repressor for gamma-herpesvirus lytic reactivation

We focused on HNRNPA2B1 for further functional analysis as its role in EBV life cycle has not been investigated. To evaluate the function of HNRNPA2B1 in EBV lytic reactivation across different cell types, we depleted HNRNPA2B1 in EBV+ B lymphoma cells and nasopharyangeal carcinoma cells.

In Akata (EBV+) Burkitt lymphoma cells, loss of HNRNPA2B1 led to a significant increase in expression of the IE protein ZTA and the early lytic protein EAD following lytic induction by IgG-crosslinking of B cell receptor (BCR), compared with cells expressing a control sgRNA (**Fig. 2A, lanes 5-6, 8-9, 11-12 vs. lanes 2-3**). Consistent with WB analysis, quantification of extracelluar EBV genome copy number further confirmed a significant increase following HNRNPA2B1 depletion (**Fig. 2B**). Consistent results were also observed in EBV+ Akata-BX1 cells, where loss of HNRNPA2B1 led to enhanced ZTA and EAD protein expression following BCR stimulation (**Fig. 2C, lanes 5-6 vs. lanes 2-3**). Akata-BX1 cells express GFP marker following lytic induction as a *GFP* gene was fused to the downstream of an early gene *BXLF1* promoter, allowing the measurement of lytic cell population by flow cytometry [35]. We found a substantial increase in the proportion of GFP+ Akata-BX1 cells upon HNRNPA2B1 depletion, indicating elevated EBV lytic reactivation at individual cell level with or without lytic induction (**Fig. 2D**) with Akata-4E3, a isogenic EBV-Akata cell line, as a control without GFP signal [36]. Conversely, overexpresison of HNRNPA2B1 in HNRNPA2B1-depleted Akata-BX1 cells suppressed IgG-crosslinking-induced ZTA and EAD protein accumulation (**Fig. 2E-2F**). This repression was accompanied by a marked decrease in GFP+ cells, further supporting that HNRNPA2B1 restricts EBV lytic replication (**Fig. 2G**).

**Figure 2.**
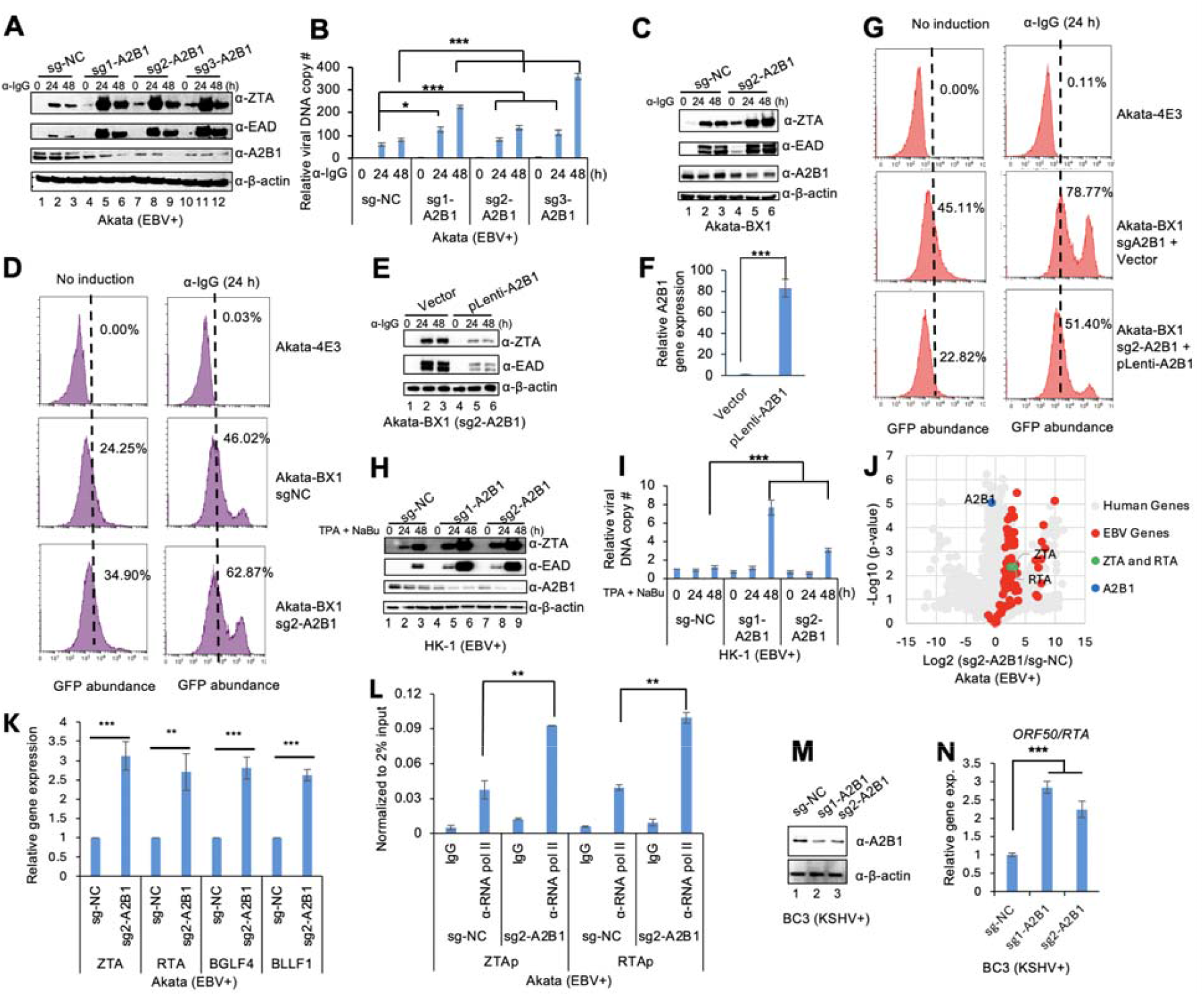
HNRNPA2B1 functions as a conserved repressor for gamma-herpesvirus lytic reactivation. (**A**) WB analysis of EBV lytic proteins ZTA and EAD in Akata (EBV+) cells expressing control sgRNA (sg-NC) or HNRNPA2B1-targeting sgRNAs following lytic induction by IgG-crosslinking for 0, 24 and 48 h. β-actin serves as a loading control. (**B**) Relative extracellular EBV DNA copy number in Akata (EBV+) cells treated as in (A) were quantified by qPCR as described in Materials and Methods. Data are presented relative to sg-NC at 0 h. Data represents ±SD from three biological replicates. *p<0.05; ***p< 0.001. (**C**) WB analysis of ZTA, EAD, and HNRNPA2B1 expression in Akata-BX1 cells expressing control sgRNA (sg-NC) or HNRNPA2B1-targeting sgRNAs following lytic induction by IgG-crosslinking. (**D**) Flow cytometric analysis of GFP reporter expression in Akata-BX1 cells carrying control sg-NC or HNRNPA2B1-targeting sgRNA (sg2-A2B1) without or with IgG-crosslinking for 24 h. EBV-Akata-4E3 cells were included as a GFP-control. (**E**) WB analysis of EBV lytic protein expression in HNRNPA2B1-depleted Akata-BX1 cells (sg2-A2B1) transduced with lentiviruses carrying empty vector or HNRNPA2B1-expressing plasmid (pLenti-A2B1) following lytic induction by IgG-crosslinking. (**F**) RT-qPCR quantification of HNRNPA2B1 transcript levels in Akata-BX1 (sg2-A2B1) cells carrying empty vector or HNRNPA2B1-expressing plasmid established in panel E. Data represents ±SD from three biological replicates. ***p< 0.001. (**G**) Flow cytometric analysis of GFP reporter expression in Akata-BX1 (sg2-A2B1) cells expressing vector control or HNRNPA2B1 without or with IgG-crosslinking for 24 h. EBV-Akata-4E3 cells were included as a GFP-control. (**H**) WB analysis of ZTA and EAD expression in HK-1 (EBV+) cells carrying control sg-NC or HNRNPA2B1-targeting sgRNAs following lytic induction by TPA and sodium butyrate (NaBu) treatment for 0, 24 and 48 h. (**I**) Relative extracellular EBV DNA copy number in HK-1 (EBV+) cells treated as in (H) was quantified by qPCR as described in Materials and Methods. Data are presented relative to sg-NC at 0 h. Data represents ±SD from three biological replicates. ***p<0.001. (**J**) Volcano plot of RNA-seq data showing differential viral gene expression in Akata (EBV+) cells expressing control sg-NC or HNRNPA2B1-targeting sgRNA (sg2-A2B1). EBV genes are shown in red; IE genes *ZTA* and *RTA* are shown in green; HNRNPA2B1 (A2B1) is shown in blue. (**K**) RT-qPCR analysis of EBV lytic gene expression in Akata (EBV+) cells expressing control sg-NC or HNRNPA2B1-targeting sgRNA (sg2-A2B1). Transcripts analyzed include *ZTA, RTA, BGLF4*, and *BLLF1*. Data are normalized to β-actin and presented as mean ± SD from three biological replicates. ** p<0.01; ***p< 0.001 (**L**) ChIP-qPCR analysis showing enrichment of RNA pol II at ZTAp and RTAp in Akata (EBV+) cells expressing control sg-NC or HNRNPA2B1-targeting sgRNA (sg2-A2B1). Data represents ±SD from three biological replicates. **p< 0.01 (**M**) WB analysis of HNRNPA2B1 expression in BC3 (KSHV+) cells expressing control sg-NC or HNRNPA2B1-targeting sgRNAs as indicated. β-actin serves as a loading control. (**N**) Relative KSHV *ORF50 (RTA)* transcript levels in BC3 (KSHV+) cells expressing control sg-NC or HNRNPA2B1-targeting sgRNAs were quantified by RT-qPCR as described in Material and Methods. Data are normalized to β-actin and presented as mean ± SD from three biological replicates. The value of sg-NC was set as 1. ***p< 0.01.

In HK-1 (EBV+) nasopharyangeal carcinoma cells [37, 38], depletion of HNRNPA2B1 enhanced ZTA and EAD protein expression following lytic induciton by TPA and sodium butyrate treatment (**Fig. 2H, lanes 5-6, 8-9 vs. lanes 2-3**), and significantly increased the extracelluar EBV genome copy number (**Fig. 2I**).

To further define the role of HNRNPA2B1 in the transcriptional regulation of EBV lytic genes, we performed global gene expression analysis in Akata (EBV+) cells carrying control sg-NC and HNRNPA2B1-targeting sgRNA (sg2-A2B1). Our RNA sequencing (RNA-seq) analysis revealed widespread upregulation of EBV lytic genes upon HNRNPA2B1 depletion, with IE genes such as *ZTA* and *RTA* among the induced transcripts (**Fig. 2J**).

To validate the transcriptomic data, we performed RT-qPCR analysis and found that the expression of IE (ZTA/RTA), early (BGLF4) and late (BLLF1) genes is significantly increased in Akata (EBV+) cells depleted of HNRNPA2B1 compared with control cells (**Fig. 2K**). We reasoned that this is due to enhanced transcriptional activity of viral genes. To test this hypothesis, we assessed RNA polymerase II (RNA Pol II) binding to EBV IE gene promoters, ZTAp and RTAp. We showed significant enrichment of RNA Pol II binding on ZTAp and RTAp relative to IgG control when HNRNPA2B1 is depleted, indicating that loss of HNRNPA2B1 promotes the transcription of EBV lytic genes (**Fig. 2L**).

To determine whether HNRNPA2B1 plays a role in the reactivation of another gamma-herpesvirus, Kaposi’s sarcoma-associated herpesvirus (KSHV), we depleted HNRNPA2B1 in KSHV+ BC3 primary effusion lymphoma cells [39]. WB analysis confirmed effective depletion of HNRNPA2B1 in BC3 (KSHV+) cells (**Fig. 2M, lanes 2-3 vs. lane 1**). Correspondingly, RT-qPCR analysis demonstrated increased expression of the KSHV IE gene *ORF50 (RTA)* following HNRNPA2B1 depletion (**Fig. 2N**). These results indicate that HNRNPA2B1 broadly suppresses the transcription of herpesvirus lytic genes.

### HNRNPA2B1 binds to EBV IE promoters

Our enChIP-MS analysis suggested that HNRNPA2B1 binds to ZTAp. HNRNPA2B1 has previously been reported to bind the herpes simplex virus 1 (HSV-1) genome during primary infection [18]. We hypothesize that HNRNPA2B1 restricts EBV lytic replication by binding to IE gene promoters, analogous to the mechanism we previously described for PIAS1 [40]. To test this hypothesis, we performed HNRNPA2B1 ChIP followed by qPCR analysis in Akata (EBV+) cells. We observed significant enrichment of HNRNPA2B1 at EBV IE promoters (ZTAp and RTAp) compared with IgG controls, with binding markedly increased at 24 h following lytic induction by IgG cross-linking (**Fig. 3A**). We reasoned that newly synthesized viral DNA may contribute to enhanced HNRNPA2B1 binding. To test this possibility, we conducted ChIP analysis in ganciclovir (GCV)-treated lytically induced Akata (EBV+) cells (**Fig. 3B-3C**). As expected, GCV treatment effectively repressed EBV DNA replication (**Fig. 3B**) [33, 41], and this repression coincided with the reduction of HNRNPA2B1 binding to the ZTAp and RTAp (**Fig. 3C**). Together, these results suggested that HNRNPA2B1 binds to EBV IE gene promoters in latency and during lytic reactivation.

**Figure 3.**
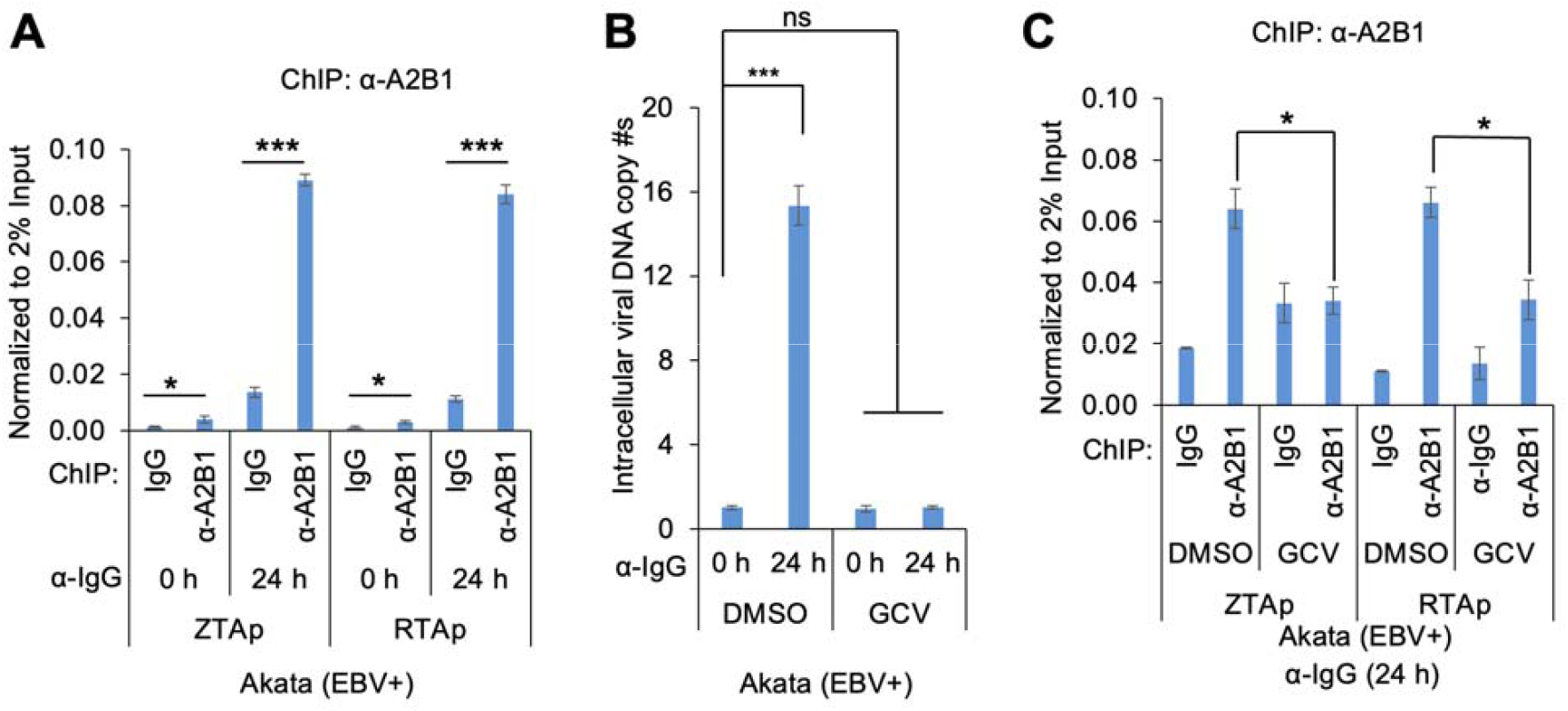
HNRNPA2B1 binds to EBV IE gene promoters. (**A**) ChIP-qPCR analysis of HNRNPA2B1 occupancy at ZTAp and RTAp in Akata (EBV+) cells following lytic induction by IgG-crosslinking for 0 h and 24 h. Anti-HNRNPA2B1 antibody was used for HNRNPA2B1 ChIP, and nonspecific IgG was used as a negative control. Data are presented as mean ± SD from three biological replicates. *p < 0.05, **p < 0.01. (**B**) Quantification of intracellular viral DNA copy number in Akata (EBV+) cells under dimethyl sulfoxide (DMSO) solvent control or GCV pretreatment followed by lytic induction for 0 h and 24 h. Viral DNA levels were measured by qPCR as described in Material and Methods. Data are presented as mean ± SD from three biological replicates. **p < 0.01; ns, not significant. (**C**) ChIP-qPCR analysis of HNRNPA2B1 enrichment at ZTAp and RTAp in lytically induced Akata (EBV+) cells pretreated with control DMSO or GCV. Anti-HNRNPA2B1 antibody was used for HNRNPA2B1 ChIP, and nonspecific IgG was used as a negative control. Data are presented as mean ± SD from three biological replicates. *p < 0.05.

### HNRNPA2B1 restricts EBV lytic reactivation by modulating H3K4 trimethylation

We demonstrated that HNRNPA2B1 binds to EBV IE gene promoters, and that HNRNPA2B1 depletion enhances RNA polymerase II occupancy at these promoters and thereby promotes the transcription of EBV IE genes. These observations prompted us to investigate whether loss of HNRNPA2B1 also affects epigenetic marks associated with transcriptional activation.

Interestingly, we found that HNRNPA2B1 depletion globally increased the activing histone mark H3K4Me3, with comparatively smaller effects on other transcription-associated marks, including H3K27ac and H2AK5ac (**Fig. 4A, lanes 4-6, 7-9, 10-12 vs lanes 1-3**). To determine whether HNRNPA2B1 directly influences H3K4Me3 level at EBV IE gene promoters, we performed ChIP qPCR analysis of H3K4Me3 occupancy at EBV ZTAp and RTAp. We found that loss of HNRNPA2B1 significantly enhanced H3K4Me3 enrichment at both promoters (**Fig. 4B**), indicating that HNRNPA2B1 normally suppresses this activating histone mark to limit EBV IE gene expression.

**Figure 4.**
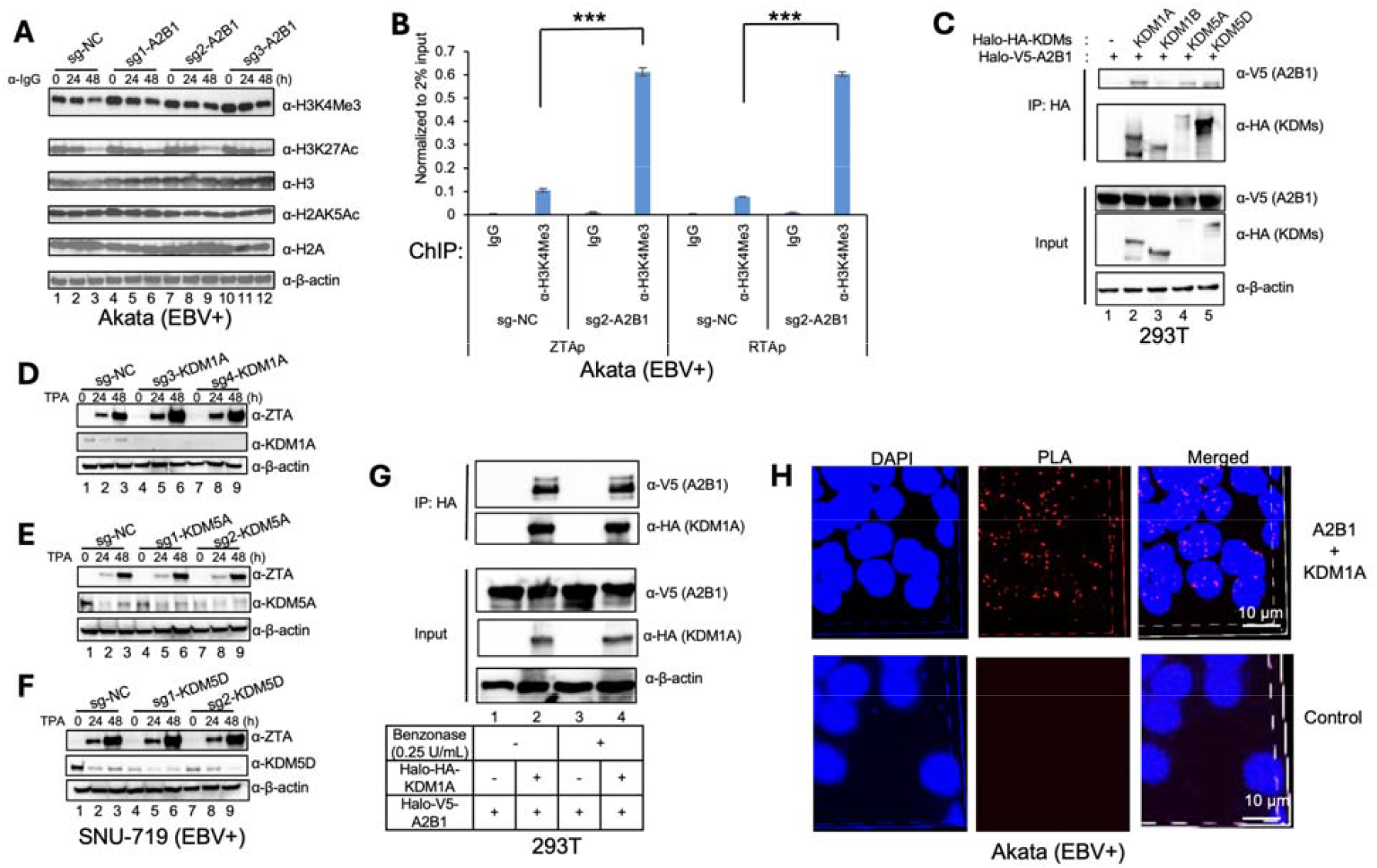
HNRNPA2B1 restricts EBV lytic reactivation by modulating H3K4Me3. (**A**) WB analysis of histone modifications in Akata (EBV+) cells following HNRNPA2B1 depletion and IgG-crosslinking-induced lytic reactivation. Whole-cell lysates were collected at the indicated time points and probed for H3K4Me3, H3K27Ac, total H3, H2AK5Ac, and total H2A. β-actin serves as a loading control. (**B**) ChIP-qPCR analysis of H3K4Me3 enrichment at EBV ZTAp and RTAp in control (sg-NC) and HNRNPA2B1-depleted (sg2-A2B1) Akata (EBV+) cells. Data are shown as relative to 2% of input. Data are presented as mean ± SD from three biological replicates. ***p<0.001. (**C**) Plasmids expressing Halo-HA-KDMs and Halo-V5-HNRNPA2B1 were co-transfected into HEK-293T cells as indicated. Cell lysates containing the indicated tagged proteins were IP-ed with anti-HA antibody-conjugated beads and analyzed by WB with anti-V5 and anti-HA antibodies. Co-IP analysis showing stronger interaction between HNRNPA2B1 and KDM1A. β-actin serves as a loading control. (**D-F**) WB analysis of EBV lytic protein ZTA and KDM1A (**D**), KDM5A (**E**), or KDM5D (**F**) expression in KDM1A-, KDM5A-, or KDM5D-depleted SNU-719 cells following lytic induction by TPA treatment. β-actin serves as a loading control. (**G**) Co-IP analysis validating the interaction between HNRNPA2B1 and KDM1A. HEK-293T cells were transfected with Halo-HA-KDM1A and Halo-V5-HNRNPA2B1 as indicated. Cell lysates were treated with or without benzonase, followed by IP using anti-HA antibody-conjugated beads. IP and input samples were analyzed by WB with anti-V5 and anti-HA antibodies. β-actin serves as a loading control. (**H**) Proximity ligation assay (PLA) demonstrating the interaction between HNRNPA2B1 and KDM1A *in situ*. Akata (EBV+) cells were blocked with 3% bovine serum albumin (BSA) in phosphate-buffered saline (PBS) for 1 h at room temperature. Subsequently, the cells were incubated with either PBS control or a combination of mouse anti-KDM1A and rabbit anti-HNRNPA2B1 antibodies. Probes were then added for ligation and amplification. Cell nuclei were visualized using Nikon AXR after staining with 4′,6-diamidino-2-phenylindole (DAPI). The interaction between HNRNPA2B1 and KDM1A *in situ* was indicated by red dot representing PLA signals. Scale bars, 10 µm.

### HNRNPA2B1 interacts with H3K4 demethylases

Because H3K4Me3 is dynamically regulated by methyltransferase and demethylases, we reasoned that HNRNPA2B1 interacts with H3K4 demethylases (KDMs) to limit H3K4Me3 level.To identify which KDMs associate with HNRNPA2B1, we cloned multiple KDM family members involved in H3K4 demethylation and assessed their interactions with HNRNPA2B1 by co-immunoprecipitation (co-IP). These experiments revealed that HNRNPA2B1 interacts KDM1A, KDM5A and KDM5D, with the strongest binding observed for KDM1A (**Fig. 4C, lane 2 vs. lanes 4-5**).

To evaluate whether these KDMs play a role in EBV reactivation, we performed CRISPR Cas9-mediated depletion of KDM1A, KDM5A, or KDM5D individually in SNU-719 cells (**Fig. 4D-4F**). We found that depletion of KDM1A resulted in a marked increase in ZTA expression following lytic induction (**Fig. 4D**), whereas knockdown of KDM5A or KDM5D had minimal effects (**Fig. 4E-4F**). These results are consistent with a prior report identifying KDM1A as a key restriction factor for EBV lytic replication [17]. Therefore, we focused on the interaction between HNRNPA2B1 and KDM1A in EBV reactivation.

To determine whether the interaction between HNRNPA2B1 and KDM1A is mediated by nucleic acids, cell lysates were treated with benzonase prior to co-IP. We observed that benzonase treatment did not affect the association between HNRNPA2B1 and KDM1A, indicating that their interaction is independent of nucleic acids (**Fig. 4G, lane 4 vs. lane 2**).

To demonstrate the interaction between HNRNPA2B1 and KDM1A in a physiologically relevant context, we performed proximity ligation assays (PLA) in Akata (EBV+) cells. Strong PLA signals localized predominantly to the nucleus were detected (**Fig. 4H**), supporting the physiological relevance of the HNRNPA2B1-KDM1A interaction.

To define the regions of KDM1A required for its interaction with HNRNPA2B1, we performed co-IP assays in HEK-293T cells transfected with plasmids expressing full-length Halo-V5-HNRNPA2B1 together with either full-length Halo-HA-KDM1A or individual Halo-HA-tagged fragments (**Fig. 5A**). Full-length KDM1A and all KDM1A fragments except the N-terminal region (amino acids 1 to 171) co-IPed with V5-HNRNPA2B1 (**Fig. 5B, lanes 2, 4, 5, and 6**). These results indicate that domains covering both SWIRM and TOWER regions of KDM1A mediate the interaction with HNRNPA2B1.

**Figure 5.**
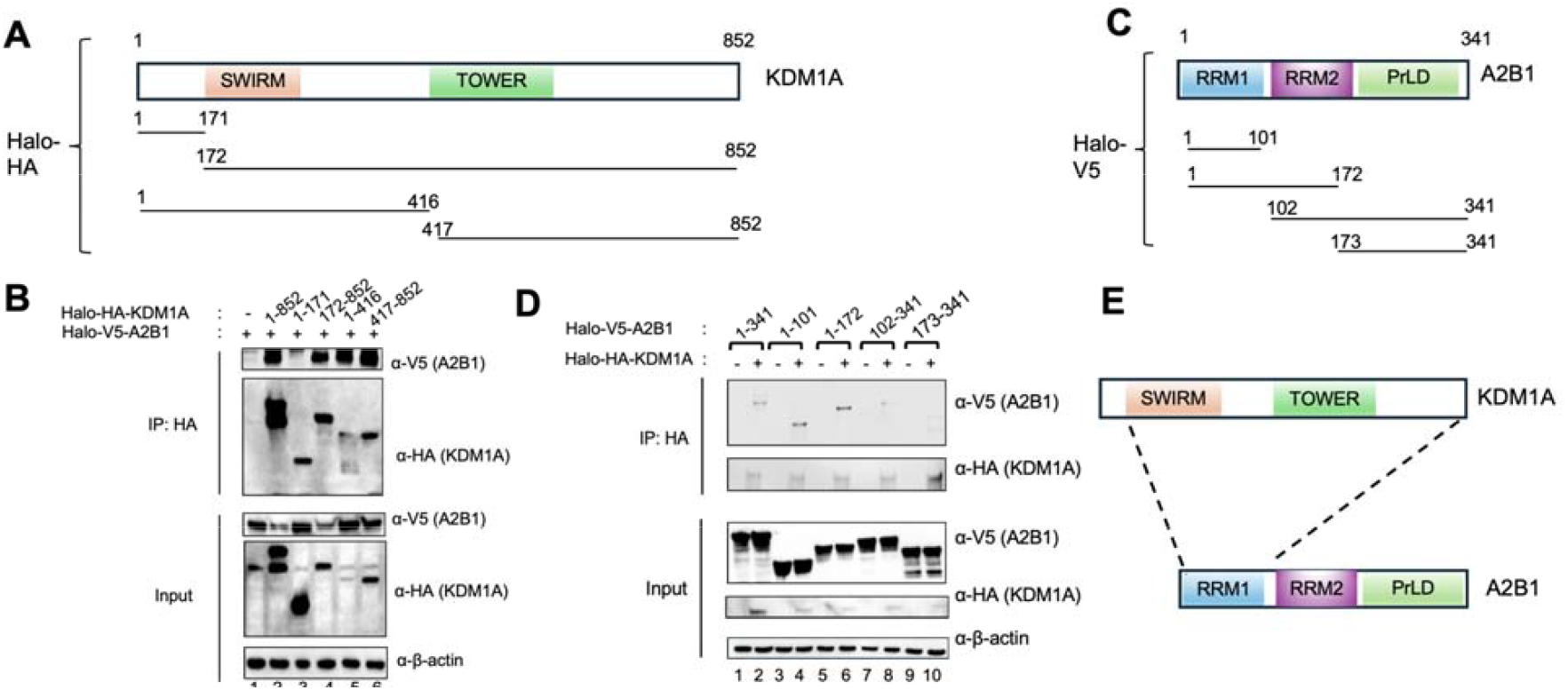
Domain-specific interaction between KDM1A and HNRNPA2B1. (**A**) Schematic representation of KDM1A domain architecture and Halo-HA-tagged KDM1A truncation constructs. SWIRM: <u>Swi</u>3p, <u>R</u>sc8p and <u>M</u>oira domain; TOWER: a pair of long, antiparallel α-helices containing domain. (**B**) HEK-293T cells were co-transfected with Halo-V5-HNRNPA2B1 and either full-length or truncated KDM1A constructs as indicated. Cell lysates containing the indicated tagged protein were IP-ed with anti-HA antibody-conjugated beads and analyzed by WB with anti-V5 and anti-HA antibodies. β-actin serves as a loading control. (**C**) Schematic representation of HNRNPA2B1 domain organization and Halo-V5-tagged HNRNPA2B1 truncation constructs. RRM: RNA recognition motif; PrLD: prion-like domain. (**D**) HEK-293T cells were co-transfected with full-length Halo-HA-KDM1A and either full-length or truncated Halo-V5-HNRNPA2B1 constructs as indicated. Cell lysates containing the indicated tagged proteins were IP-ed with anti-HA antibody-conjugated beads and analyzed by WB with anti-V5 and anti-HA antibodies. β-actin serves as a loading control. (**E**) Proposed model illustrating the domains required for the interaction between KDM1A and HNRNPA2B1.

To determine the regions of HNRNPA2B1 involved in KDM1A binding, HEK-293T cells were co-transfected with plasmids expressing full-length Halo-HA-KDM1A and either full-length Halo-V5-HNRNPA2B1 or individual Halo-V5-tagged fragments (**Fig. 5C**). Immunoprecipitation of KDM1A using anti-HA antibody-conjugated beads demonstrated that the N-terminal regions of HNRNPA2B1 [amino acids (aa) 1-101 or 1-172] were strongly co-IPed with HA-KDM1A, whereas the C-terminal regions (aa 102-341 or 173-341) showed much weaker association (**Fig. 5D, lanes 4 and 6 vs. lane 8 and 10**). Together, these data suggest that the C-terminal domain of KDM1A interacts specifically with the RNA recognition motif 1 (RRM1) of HNRNPA2B1 (**Fig. 5E**).

### HNRNPA2B1 recruits KDM1A to EBV IE promoters

We hypothesize that HNRNPA2B1 recruits KDM1A to EBV IE gene promoters to limit H3K4Me3 level. To test this hypothesis, we performed ChIP-qPCR analysis of KDM1A enrichment at ZTAp and RTAp in control and HNRNPA2B1-depleted Akata (EBV+) cells. We found that loss of HNRNPA2B1 resulted in a significant reduction of KDM1A occupancy at ZTAp and RTAp (**Fig. 6A**). Because KDM1A expression was not affected by HNRNPA2B1 depletion (**Fig. 6B, lanes 4-6 vs lanes 1-3**), the reduced KDM1A promoter occupancy in HNRNPA2B1-depleted cells reflects impaired recruitment rather than KDM1A abundance.

**Figure 6.**
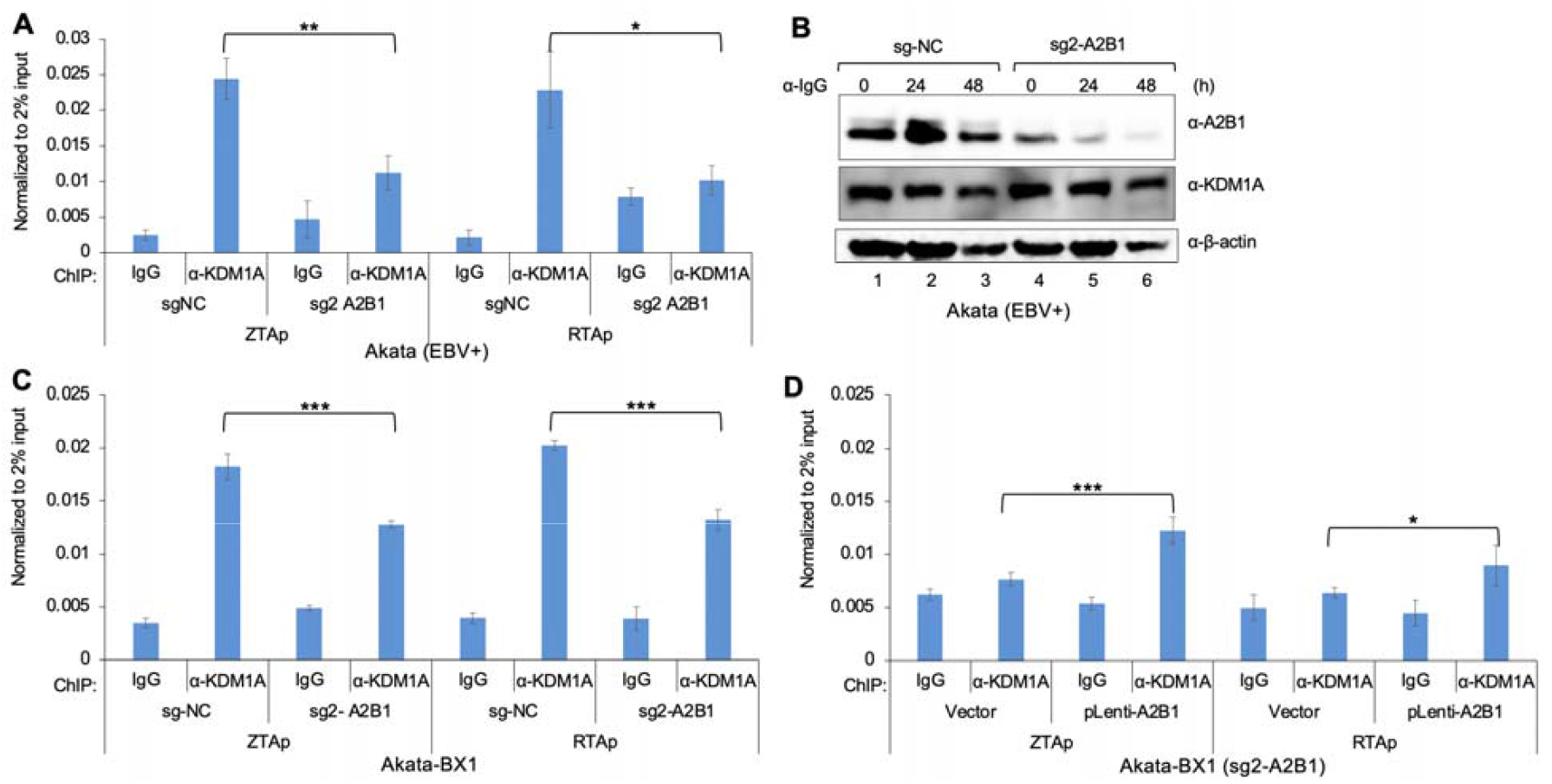
HNRNPA2B1 promotes KDM1A recruitment to EBV IE gene promoters. (**A**) ChIP-qPCR analysis of KDM1A occupancy at ZTAp and RTAp in Akata (EBV+) cells transduced with non-targeting control (sg-NC) or HNRNPA2B1-targeting sgRNA (sg2-A2B1). Anti-KDM1A was used for KDM1A ChIP and IgG served as a negative control. Data are normalized to 2% input and shown as mean ± SD from three biological replicates. *p<0.05; **p<0.01. (**B**) Akata (EBV+) cells carrying control sg-NC or HNRNPA2B1-targeting sgRNA (sg2-A2B1) were treated with anti-human IgG for the indicated times. Protein levels of HNRNPA2B1 and KDM1A were analyzed by WB. β-actin serves as a loading control. (**C**) ChIP-qPCR analysis of KDM1A occupancy at ZTAp and RTAp in Akata-BX1 cells expressing control sg-NC or HNRNPA2B1-targeting sgRNA (sg2-A2B1). Anti-KDM1A was used for KDM1A ChIP and IgG served as a negative control. Data are normalized to 2% input and shown as mean ± SD from three biological replicates. ***p<0.001. (**D**) ChIP-qPCR analysis of KDM1A occupancy at ZTAp and RTAp in Akata-BX1 sgA2B1 cells reconstituted with vector control or HNRNPA2B1 (pLenti-A2B1). Anti-KDM1A antibody was used for KDM1A ChIP and IgG served as a negative control. Data are normalized to 2% input and shown as mean ± SD from three biological replicates. *p<0.05; ***p<0.001.

Consistent with the findings in Akata (EBV+) cells, depletion of HNRNPA2B1 in Akata-BX1 cells led to reduced KDM1A binding at both ZTAp and RTAp (**Fig. 6C**). To further confirm these results, we restored HNRNPA2B1 expression in HNRNPA2B1-depleted Akata-BX1 cells. We noticed that HNRNPA2B1 restoration resulted in enhanced recruitment of KDM1A to ZTAp and RTAp, as determined by KDM1A ChIP-qPCR analysis (**Fig. 6D**). Together, these results demonstrate that HNRNPA2B1 fosters the recruitment of KDM1A to EBV ZTAp and RTAp, thereby contributing to the maintenance of a repressive chromatin state to maintain vial latency and restrict lytic reactivation.

## DISCUSSION

EBV latency is maintained by stringent repression of IE gene expression, yet the host factors that promote this silencing at the viral chromatin level remain incompletely defined [5-7]. Using enChIP-MS, we directly captured host proteins associated with EBV ZTAp in its native chromatin context, facilitating unbiased identification of putative factors involved in *ZTA* gene repression. This approach has previously been used to identify the nucleosome remodeling and deacetylase complex at D4Z4 repeat in myoblasts, host proteins associated with the parvovirus B19 genome, and EPAS1 promoter-associated proteins in neuroblastoma cells, supporting its utility for locus-specific chromatin proteomics [30-32]. Among approximately 1000 proteins identified, we focused on HNRNPs for further functional analysis and discovered a previously unappreciated role for HNRNPA2B1 in EBV latency and reactivation.

HNRNPA2B1 has been reported as an anti-viral or pro-viral factor in different context. For example, HNRNPA2B1 restricts hepatitis B virus (HBV) and SARS-CoV2 replication by triggering anti-viral immune response via TBK1-IRF3 pathway [21]. During HSV-1 infection, HNRNPA2B1 promotes interferon production via binding to viral DNA and promoting the trafficking of *CGAS, IFI16*, and *STING* mRNAs, thereby restricting viral replication [18]. In contrast, overexpression of HNRNPA2B1 in THP-1 cells has been shown to enhance gene expression for several viruses, including severe fever with thrombocytopenia syndrome virus (SFTSV) [26]. In Senecavirus A model, the interaction between viral VP3 protein and HNRNPA2B1 facilitates viral internal ribosome entry site (IRES)-driven protein translation while simultaneously suppressing host cell translation and interferon response [22].

In our study, we demonstrated that HNRNPA2B1 consistently suppresses EBV lytic reactivation across multiple EBV+ cell models. Intriguingly, we found that depletion of HNRNPA2B1 enhances expression of IE and downstream lytic genes, increases RNA polymerase II occupancy at EBV IE promoters, increases the frequency of cells entering the lytic cycle, and promotes the production of extracellular viral genomes. These effects were observed in EBV+ cancer models derived from both B cells and epithelial cells, indicating that HNRNPA2B1-mediated viral gene repression is not cell type specific. Conversely, ectopic expression of HNRNPA2B1 suppressed lytic replication, supporting a direct role for HNRNPA2B1 in maintaining viral latency.

One novel aspect from our study is the establishment of HNRNPA2B1 as a regulator of viral chromatin accessibility. Loss of HNRNPA2B1 led to increased enrichment of activating histone marks, particularly H3K4me3, at the IE promoters, consistent with enhanced transcriptional activity. We further showed that HNRNPA2B1 interacts with the histone demethylase KDM1A in a nucleic acid-independent manner and is required for efficient recruitment of KDM1A to EBV IE promoters. Disruption of this pathway reduced KDM1A occupancy, promoted accumulation of H3K4Me3 mark, and facilitated lytic transcription.

KDM1A, also known as LSD1, functions canonically as part of the CoREST corepressor complex. Recent work has shown that recruitment of KDM1A to the ZTA promoter depends on the transcription factor ZNF217 to regulate H3K4 methylation level [17]. Whether HNRNPA2B1 is a component of a CoREST-KDM1A-ZNF217 regulatory complex at EBV IE promoters remains to be determined.

HNRNPA2B1 plays a pivotal role in a variety of cellular processes, particularly RNA metabolism [44, 45]. While HNRNPA2B1 is primarily recognized for its RNA-binding capabilities, our findings together with recent studies suggest that HNRNPA2B1 may also function as a chromatin-associated factor to control gene expression [27, 28, 46]. Our findings expand this functional repertoire by demonstrating a direct role of HNRNPA2B1 in chromatin-based transcriptional repression of a latent viral genome. The observation that HNRNPA2B1 associates with IE promoters and regulates RNA polymerase II recruitment suggests a coordinated control of transcription initiation and early RNA processing by HNRNPA2B1.

In summary, this study identifies HNRNPA2B1 as a previously unrecognized epigenetic regulator that restricts EBV lytic reactivation through KDM1A-dependent histone H3K4 demethylation at IE promoters (**Fig. 7**). These findings provide new insights into how host RNA-binding proteins interact with histone demethylase to control herpesvirus gene expression and highlight a latency control mechanism that may be broadly relevant to other latent DNA viruses.

**Figure 7.**
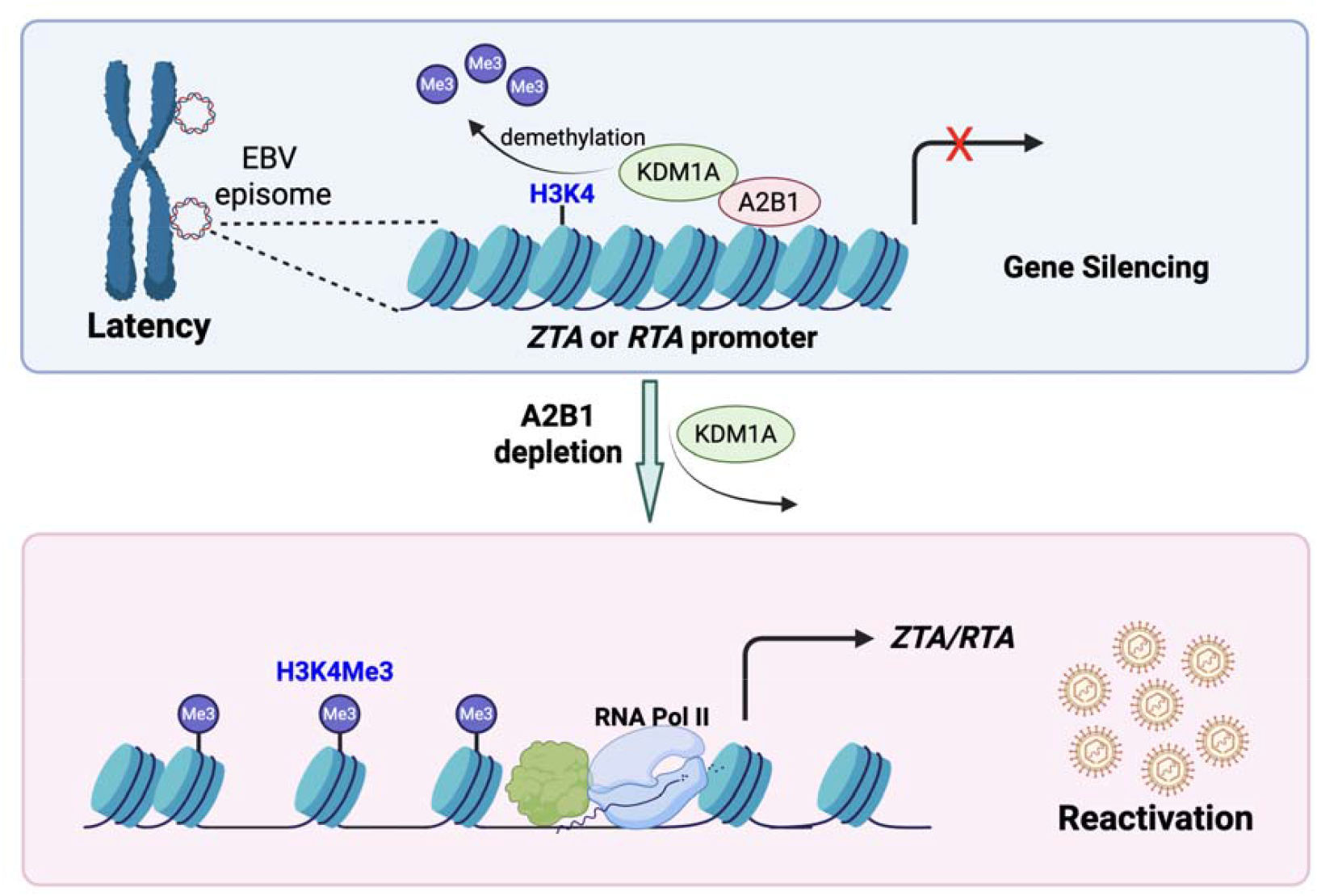
Model summarizing the role of HNRNPA2B1 in promoting EBV latency and suppressing lytic reactivation. HNRNPA2B1 (A2B1) recruits KDM1A to EBV *ZTA* and *RTA* promoters, leading to H3K4 demethylation and repression of lytic gene expression, thereby maintaining viral latency. Depletion of HNRNPA2B1 diminishes KDM1A recruitment to EBV IE gene promoters, increases H3K4Me3 levels, and enhances RNA polymerase II occupancy, resulting in activation of EBV lytic gene expression and viral reactivation.

## MATERIAL AND METHODS

### Cell lines and cultures

Akata (EBV+), SNU-719 (EBV+), HK-1 (EBV+), Akata-BX1 (EBV+), and BC3 (KSHV+) cells were cultured in Roswell Park Memorial Institute medium (RPMI 1640) supplemented with 10% FBS (Cat. # 26140079, Thermo Fisher Scientific) in 5% CO_2_ at 37°C [40, 47-50]. HK-1 (EBV+) cells (courtesy of Dr. George Tsao, Hong Kong University) were supplemented with 800 μg/mL G418 in the culture medium [33, 37, 38, 51]. Akata-BX1 (EBV+) cells were supplemented with 495 μg/mL G418 in the culture medium [35]. HEK-293T cells were cultured in Dulbecco’s modified eagle medium (DMEM) supplemented with 10% FBS in 5% CO_2_ at 37°C [52, 53]. See also **Table 1** for cell line sources.

**Table 1.** Antibody, reagents, constructs and cell lines.

| REAGENT<br>RESOURCE | OR<br>SOURCE | IDENTIFIER |
| --- | --- | --- |
| **Antibodies and reagents** |  |  |
| Mouse Anti-HNRNPA2B1 | Proteintech | Cat #67445-1-Ig |
| Rabbit Anti-HNRNPA2B1 | Proteintech | Cat #14813-1-AP |
| Rabbit Anti-HNRNPAB | Proteintech | Cat #31175-1-AP |
| Rabbit Anti-HNRNPF | Proteintech | Cat #14974-1-AP |
| Rabbit Anti-HNRNPG | Cell Signaling Technology | Cat #14794T |
| Rabbit Anti-HNRNPI | Proteintech | Cat #12582-1-AP |
| Rabbit Anti-HNRNPM | Proteintech | Cat #26897-1-AP |
| Rabbit Anti-HNRNPP2 | Proteintech | Cat #11570-1-AP |
| Rabbit Anti-HNRNPR | Proteintech | Cat #29980-1-AP |
| Mouse Anti-ZTA(BZ1) | Santa Cruz | Cat #sc-53904 |
| Mouse Anti-RNA Polymerase II | Milipore Sigma | Cat #05-623-25UG |
| Mouse IgG | Santa Cruz | Cat. #sc-2025 |
| Rabbit Anti-EBV EA-D-p52/50 Antibody, clone R3 | Milipore Sigma | Cat #MAB8186 |
| Mouse anti-β-actin antibody | Santa Cruz | Cat #sc-47778 |
| Anti-V5-HRP | Thermo Fisher | Cat #R960-25 |
| Anti-HA | Roche | Cat #11-867-431-001 |
| Rabbit-Anti H3K4Me3 | Cell Signaling Technology | Cat # 9751S |
| Acetyl-Histone Antibody Sampler Kit | Cell Signaling Technology | Cat #9933 |
| JARID1/KDM5 Histone Demethylase Antibody Sampler Kit | Cell Signaling Technology | Cat #25497T |
| Rabbit Anti-H3K27Ac | Cell Signaling Technology | Cat #4353S |
| Rabbit Anti-H3 (D2B12) | Cell Signaling Technology | Cat #4620S |
| Rabbit Anti-KDM1A | Cell Signaling Technology | Cat #2139S |
| Mouse Anti-KDM1A | Proteintech | Cat #67037-1-Ig |
| ChromoTek DYKDDDDK Fab-Trap® Agarose beads | Proteintech | Cat #ffa |
| Goat Anti-Human IgG (whole molecule) | MP Biomedicals | Cat #0855087 |
| Phorbol-12-myristate-13-acetate | Milipore Sigma | Cat #52-440-01MG |
| Sodium Butyrate | TCI America | Cat #S051925G |
| PEI Max | Polyscience | Cat #24765-100 |
| Anti-V5 magnetic beads | MBL | Cat #M167-11 |
| Anti-HA magnetic beads | Thermo Fisher | Cat #88836 |
| SimpleChIP® Enzymatic | Cell Signaling Technology | Cat #9003S |
| Chromatin IP Kit (Magnetic Beads) |  |  |
| RQ1 RNase-free DNase | Promega | Cat #M6101 |
| Proteinase K | Meridian Bioscience | Cat #BIO-37084 |
| ISOLATE II RNA Mini Kit | Meridian Bioscience | Cat #BIO-52073 |
| High-Capacity cDNA Reverse Transcription Kit with Rnase Inhibitor | Applied Biosystem | Cat #4374966 |
| NaveniFlex Cell Red Kit | Navinci | Cat #NC.MR.100 |
| Ganciclovir | Milipore Sigma | Cat #PHR1593 |
| **Constructs** |  |  |
| pMD2.G | Addgene | plasmid #12259 |
| psPAX2 | Addgene | plasmid #12260 |
| pHTN-V5-HNRNPA2B1 | This study | pFS80 |
| pLenti-HNRNPA2B1 | This study | pFS112 |
| pHTN-HA-KDM1A | This study | pFS574 |
| pHTN-HA-KDM1B | This study | pFS688 |
| pHTN-HA-KDM5A | This study | pFS689 |
| pHTN-HA-KDM5D | This study | pFS687 |
| **Cell lines** |  |  |
| Akata (EBV+) | Hayward Lab Collection | NA |
| SNU-719 | Hayward Lab Collection | NA |
| Akata-BX1 | Hayward Lab Collection | NA |
| 293T cells | Hayward Lab Collection | NA |
| BC3 | Hayward Lab Collection | NA |
| HK-1 (EBV+) | [37, 38, 51] | NA |

### Plasmid construction

The non-targeting control (sg-NC) and ZTAp-targeting sg5 oligonucleotides from our previous study [33] were cloned into pZLCv2-3xFLAG-dCas9-HA-2xNLS vector (Cat. #106357, Addgene; gift from Stephen Tapscott) using methods as previously described [30]. HNRNPA2B1, KDM1A, KDM5A, KDM5D, and KDM1B coding sequences were amplified by PCR using Q5 High-Fidelity polymerase from the following templates: MBP-hnRNPA2_FL_WT (Cat. #98662, Addgene; gift from Nicolas Fawzi), pIDS-LSD1fl (Cat. #109157, Addgene; gift from Monika Golas), TFORF2771-KDM5A (Cat. #143614, Addgene; gift from Feng Zhang), TFORF2768-KDM5D (Cat. #142159, Addgene; gift from Feng Zhang), and pLenti6.3/V5-DEST-KDM1B (Cat. HsCD00963160, DNAsu). The PCR products were cloned into the pHTN-CMV-Neo vector. HNRNPA2B1 was tagged with V5 at the N-terminus, while all KDM constructs carried an N-terminal HA tag. HNRNPA2B1, KDM1A, and KDM1B were contructed using the Gibson assembly method, whereas KDM5A and KDM5D were inserted into the vector following EcoRI/NotI double digestion and DNA ligation. All primer sequences are listed in **Table 2**.

**Table 2.**
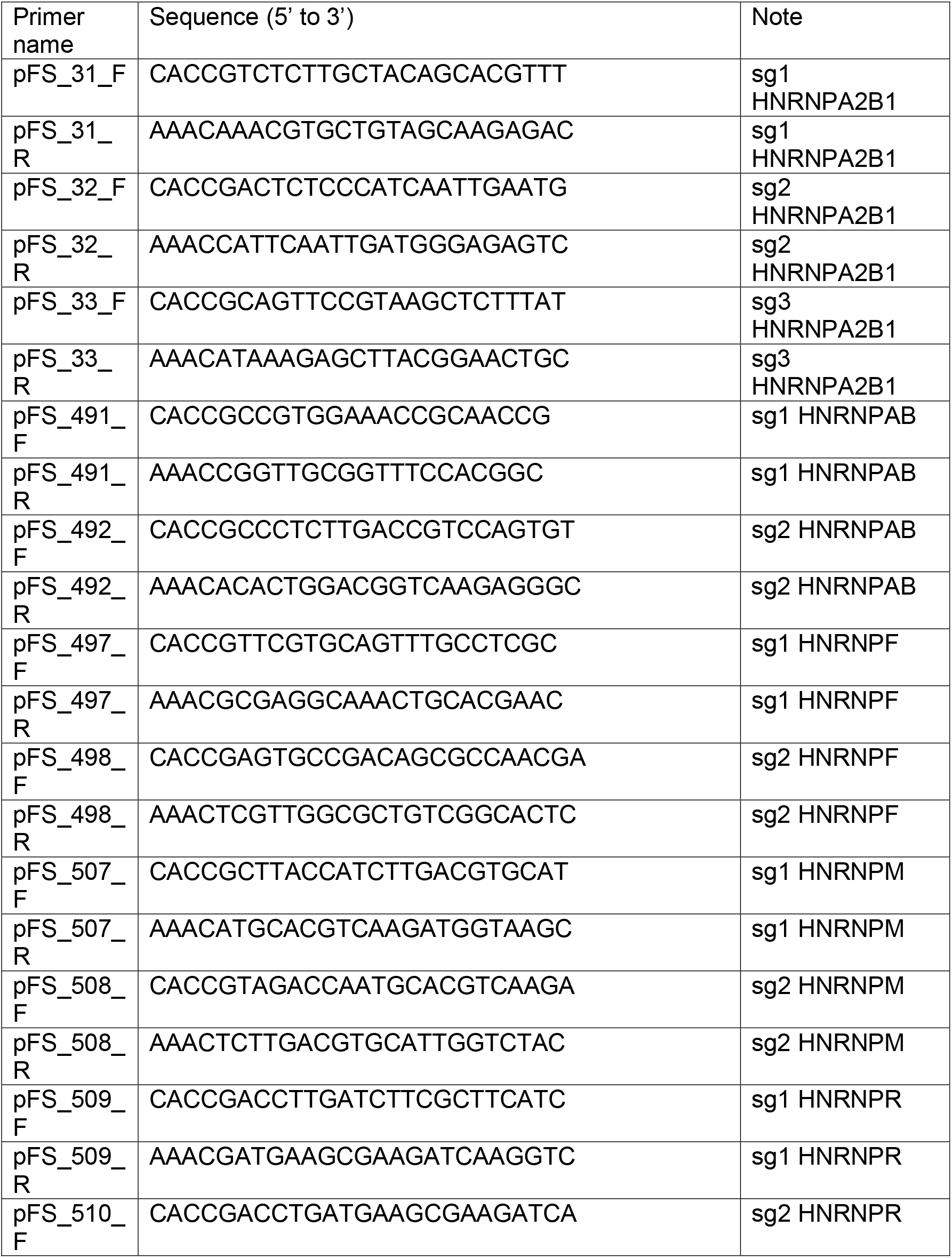
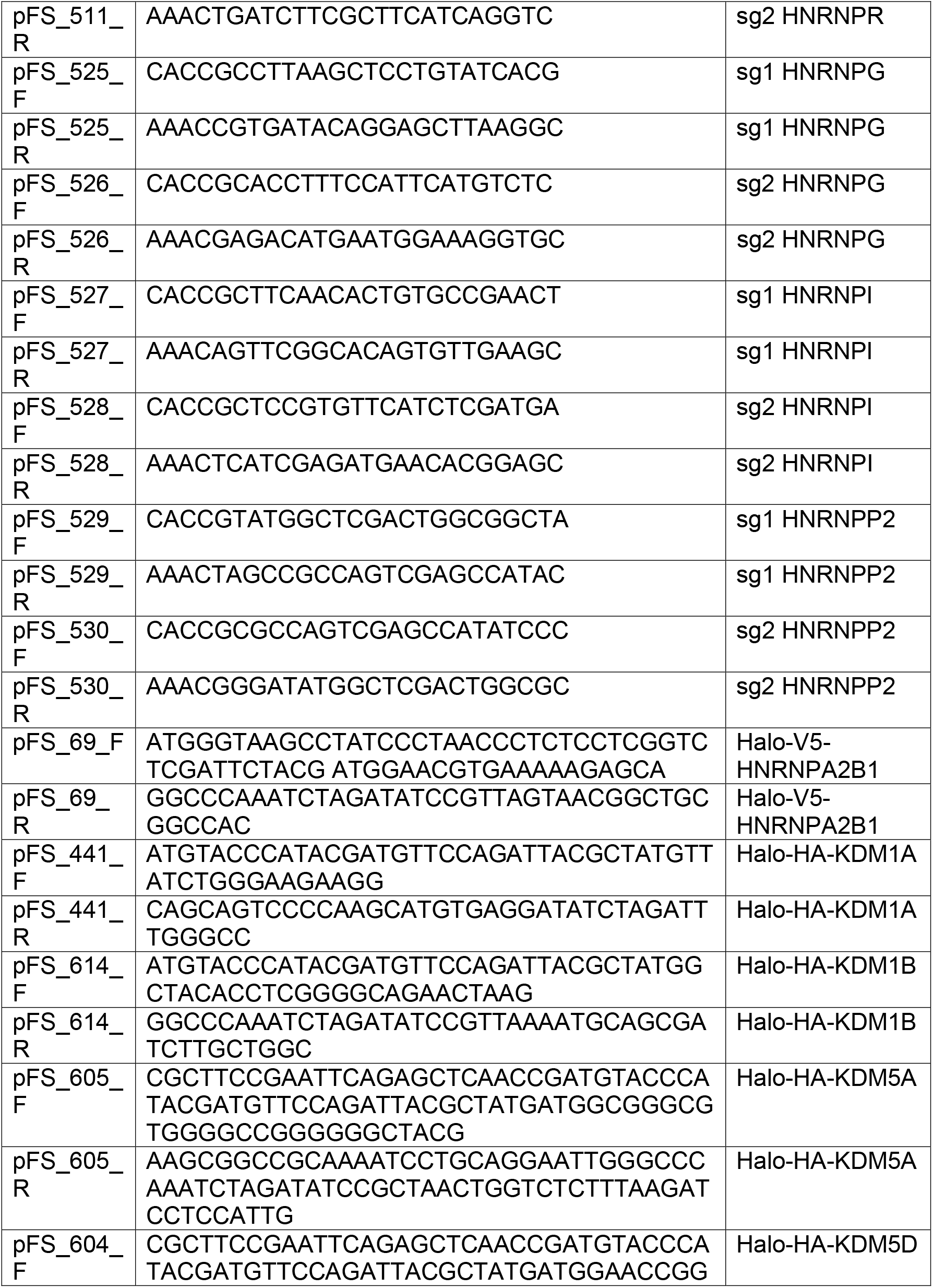
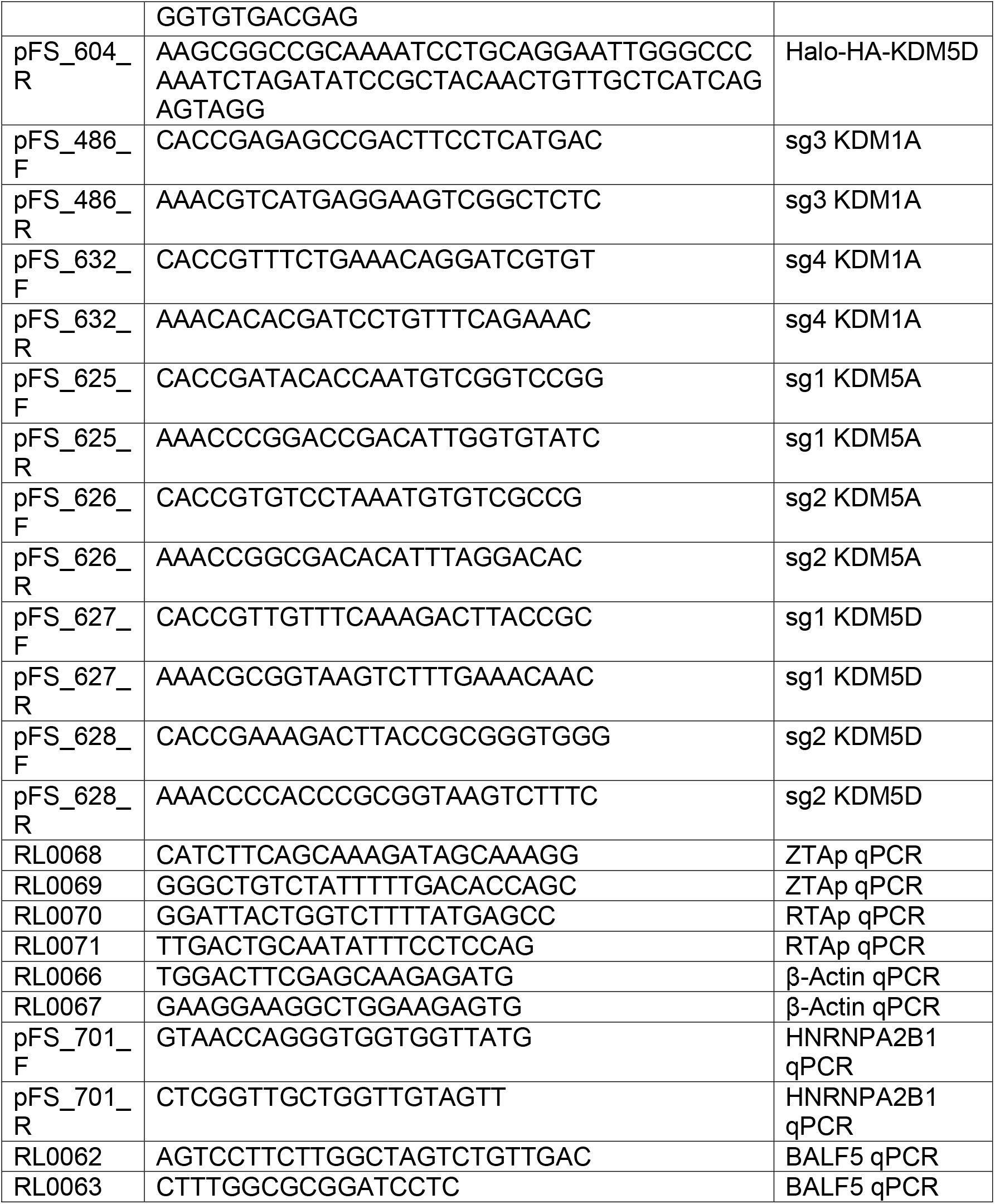
Primer List.

### Generation of stable cell line

Lentiviruses isolated from the HEK293-T medium were used to infect the Akata (EBV+), HK-1 (EBV+) and SNU-719 (EBV+) cells. Forty-eight hours post-transduction, the cells were cultured in the presence of puromycin (2 µg/mL) or blasticidin (10 μg/mL) for cell line establishment.

### enChIP-MS

enChIP-MS was modified from a previously described protocol [30]. 4 × 10^7^ cells were collected by centrifugation at 1000 rpm for 5 minutes at room temperature using a swinging bucket rotor. The cell pellet was resuspended in 1 mL of ice-cold Cell Lysis Buffer (10 mM Tris-HCl pH 8.0, 10 mM NaCl, 0.2% NP-40, EDTA-free protease inhibitor and PMSF) and incubated on ice for 10 minutes to release nuclei.

Nuclei were pelleted at 2500 rpm for 5 minutes at 4 °C in a swinging bucket centrifuge and resuspended in 37 mL room temperature PBS. Crosslinking was performed by adding 1.5 mL of 37% formaldehyde to achieve a final concentration of 1.5%, followed by gentle rocking for 15 minutes at room temperature. The reaction was quenched by adding 4 mL of 10 × glycine (Cat. #9003S, Cell Signaling Technology) and incubating for an additional 5 minutes at room temperature with rocking. Crosslinked nuclei were pelleted at 2000 rpm for 10 minutes at 4 °C, resuspended in 1 mL cold PBS, transferred to a 1.5 mL tube, and centrifuged again at 2000 rpm for 5 minutes at 4 °C.

The supernatant was removed and the pellet was washed with 2 mL ice cold 1 × Buffer B (Cat. #9003S, Cell Signaling Technology) supplemented with 1 µL DTT, followed by centrifugation and removal of the supernatant.

The nuclei were then resuspended in 200 µL 1 × Buffer B containing 0.1 µL DTT and transferred to a 1.5 mL microcentrifuge tube. Micrococcal nuclease digestion was initiated by adding 5 µL enzyme, mixing gently by inversion, and incubating at 37 °C for 20 minutes with mixing every 3 minutes. The reaction was stopped by adding 10 µL EDTA and placing the tube on ice for 2 minutes. Nuclei were pelleted at 16,000 × g for 1 minute at 4 °C and the supernatant was discarded.

The nuclear pellet was resuspended in 200 µL 1 × ChIP Buffer (Cat. #9003S, Cell Signaling Technology) supplemented with 1 µL of 200 × protease inhibitor cocktail (Cat. #9003S, Cell Signaling Technology) and incubated on ice for 10 minutes. Samples were sonicated using 10 second pulse on and 10 second pulse off cycles for five rounds, with 30 seconds incubation on ice between rounds. Lysates were clarified by centrifugation at 9,400 × g for 10 minutes at 4 °C.

For immunoprecipitation, we utilized ChromoTek DYKDDDDK Fab-Trap Agarose beads (Cat. #ffa, Proteintech). Briefly, 160 µL FLAG-Trap beads were equilibrated in 2 mL IP Wash I buffer (10 mM Tris-HCl pH 7.5, 150 mM NaCl, and 0.25% NP-40) and centrifuged at 2,500 × g for 2 minutes at 4 °C. The wash was repeated once.

The clarified chromatin lysate was diluted with 300 µL 1 × ChIP Buffer supplemented with 1.5 µL 200 × protease inhibitor cocktail and incubated overnight at 4 °C with the prepared beads. Beads were collected by centrifugation at 2,500 × g for 2 minutes at 4 °C and washed twice with 2 mL ice cold IP Wash I buffer.

Beads were subsequently washed three times with 2 mL ice cold IP Wash II buffer (10 mM Tris-HCl pH 7.5 and 150 mM NaCl), each followed by centrifugation at 2,500 × g for 2 minutes at 4 °C. After the final wash, beads were resuspended in 160 µL ice cold IP Wash II buffer, divided equally into two tubes of approximately 80 µL each, centrifuged again at 2,500 × g for 2 minutes at 4 °C, and the supernatant was removed. Samples were stored at −80 °C for downstream MS analysis by the Taplin Mass Spectrometry Facility at Harvard Medical School.

Beads were washed at least five times with 100 µl of 50 mM ammonium bicarbonate. Modified sequencing-grade trypsin (Promega, Madison, WI) was added at 5 µl (200 ng/µl), and samples were incubated at 37 °C overnight for proteolytic digestion. Following digestion, samples were centrifuged or placed on a magnetic rack, depending on bead type, and the supernatant was collected. Peptide extracts were dried using a SpeedVac concentrator for approximately 1 h, resuspended in 50 µl of HPLC solvent A (2.5% acetonitrile, 0.1% formic acid), and desalted using STAGE tips [54].

On the day of analysis, samples were reconstituted in 10 µl of HPLC solvent A. Peptides were separated using a nano-scale reverse-phase HPLC system equipped with a capillary column packed in-house with 2.6 µm C18 spherical silica beads in a fused silica capillary (100 µm inner diameter, approximately 30 cm in length) with a flame-drawn tip [55]. After column equilibration, samples were loaded using a Famos autosampler (LC Packings, San Francisco, CA). Peptides were eluted using a linear gradient of increasing concentrations of solvent B (97.5% acetonitrile, 0.1% formic acid).

Eluting peptides were ionized by electrospray ionization and analyzed on a Velos Orbitrap Elite ion trap mass spectrometer (Thermo Fisher Scientific, Waltham, MA). Peptides were detected, isolated, and fragmented to generate tandem mass spectra. Peptide sequences and corresponding protein identities were assigned by database searching of the acquired spectra using Sequest (Thermo Fisher Scientific, Waltham, MA) [56]. Databases included reversed sequence entries to estimate false discovery rates, and peptide identifications were filtered to achieve a false discovery rate of 1 to 2 percent (**see Table S1**).

### ChIP-qPCR

ChIP process and qPCR were performed as previously described [57]. DNA-protein complex were immunoprecipitated with anti-RNA polymerase II (Cat. #05-623-25UG, Milipore Sigma), anti-HNRNPA2B1 (Cat. # 67445-1-Ig, Proteintech), anti-H3K4Me3 (Cat. # 9751S, Cell Signaling Technology), anti-KDM1A (Cat. #2139S, Cell Signaling Technology), rabbit IgG control (Cat. #2729, Cell Signaling Technology), and mouse IgG control (Cat. #sc-2025, Santa Cruz). ChIP was performed using an SimpleChIP Enzymatic Chromatin IP kit (Cat. #9003S, Cell Signaling Technology) as described previously [40]. ChIP-ed DNA was quantified by qPCR using *ZTA* and *RTA* primers (**Table 2**).

### Cell lysis, WB and immunoprecipitation

Cell lysis, WB and immunoprecipitation (IP) were performed as previously described [58], with minor modifications. Cells were harvested, lysed in 2 × SDS-PAGE sample buffer, and boiled for 5 minutes. Proteins were separated on 4-20% TGX gels (Cat. #4561096; Bio-Rad), transferred to PVDF membranes, and probed with the indicated primary antibodies followed by horseradish peroxidase-conjugated secondary antibodies. See also **Table 1** for antibody sources.

For IP, cells were lysed on ice for 30 minutes in buffer containing 50 mM Tris-HCl (pH 7.5), 150 mM NaCl, 0.1% NP-40, and protease inhibitor cocktail (Cat. #4693116001; Sigma-Aldrich). Lysates were sonicated (10 seconds on/10 seconds off, three cycles, 35% amplitude) and clarified by centrifugation at 14,600 × g for 15 minutes at 4 °C. Ten percent of the supernatant was reserved as input, and the remainder was incubated with the indicated magnetic beads. Input and IP-ed proteins were analyzed by WB using the specified antibodies.

### Flow cytometry

Approximately 5 × 10^6^ cells were harvested and maintained on ice, washed once with cold PBS, and pelleted by centrifugation at 1,200 × g for 5 minutes. Cells were resuspended in PBS and fixed with 4% paraformaldehyde for 10 minutes at room temperature, followed by two washes with PBS and resuspension in PBS. Immediately prior to data acquisition, samples were passed through a 40 µm cell strainer to remove aggregates. Fluorescence signals were acquired on a CytoFLEX 2-L flow cytometer and detected using a 525 nm filter (Beckman-Coulter). Data were analyzed by CytExpert 2.6 software (Beckman-Coulter).

### Lytic induction and EBV copy number detection

Akata (EBV+) cells were seeded at 1 × 10□ cells/mL in 6-well plates. After 3 h, cells were lytically induced by anti-human IgG (50 µg/mL; Cat. #0855087, MP Biomedicals) and harvested at the indicated time points as described previously [2, 40, 58]. SNU-719 (EBV+) and HK-1 (EBV+) cells were seeded at 3 × 10□ cells/mL in 6-well plates and incubated overnight. On the following day, SNU-719 cells were treated with TPA (20 ng/mL; Cat. #NC9325685, Fisher Scientific), while HK-1 cells were treated with a combination of TPA (40 ng/mL) and sodium butyrate (5 mM; Cat. #S0519, Tokyo Chemical Industry) to induce EBV lytic cycle.

To quantify EBV replication, intracellular EBV DNA and virion-associated DNA were measured by qPCR. For intracellular EBV DNA analysis, total genomic DNA was extracted using a genomic DNA purification kit (Cat. # A1120, Promega) according to the manufacturer’s instructions. Briefly, 2 × 10^6^ cells were harvested and resuspended in Nuclei Lysis Solution, followed by RNA removal by RNase A treatment. Proteins were precipitated with Protein Precipitation Solution, and the clarified supernatant was mixed with isopropanol to precipitate genomic DNA. The DNA pellet was washed with ethanol and resuspended in DNA Rehydration Solution. EBV DNA levels were quantified using primers targeting *BALF5* and normalized to β*-actin* genomic DNA.

Extracellular virion-associated DNA was extracted and quantified following established protocols [58, 59]. Briefly, 120 µl EBV-containing culture supernatants were treated with RQ1 RNase-free DNase (Cat. #M6101; Promega) to remove free-floating DNA. The reaction was stopped using the supplied stop buffer, and then Proteinase K (Cat. #BIO-37084; Meridian Bioscience) and SDS were added to digest viral proteins and release virion-associated DNA. DNA was purified by phenol-chloroform extraction and precipitated with 250 μL of 100% isopropanol, 50 μL of 3M sodium acetate pH 5.2, and 1 μL glycogen at −80 °C overnight. Pellets were washed with 70% ethanol, air-dried, and resuspended in Tris-EDTA (TE) buffer (10 mM Tris, 1 mM EDTA, pH 8.0). EBV DNA was quantified by qPCR using *BALF5*-specific primers [58].

### Ganciclovir treatment

Akata (EBV+) cells were pretreated with ganciclovir (GCV; 10 µg/mL) or solvent dimethyl sulfoxide (DMSO) as a control [33] for 1 h prior to anti-human IgG-mediated crosslinking of BCR to induce EBV lytic replication.

### RNA isolation, cDNA production, and RNA-seq

Total RNA was isolated using the ISOLATE II RNA Mini Kit (Merdian Bioscience) according to the manufacturer’s instructions. Briefly, cell pellets containing up to 5 × 10^6^ cells were lysed in RLY buffer supplemented with β-mercaptoethanol and homogenized by vortexing. Lysates were clarified using ISOLATE II filters, and RNA binding conditions were adjusted by the addition of 70% ethanol. Samples were loaded onto silica spin columns, followed by membrane desalting and on-column DNase I treatment to remove genomic DNA. Columns were sequentially washed with RW1 and RW2 buffers, dried by centrifugation, and total RNA was eluted in RNase-free water. For gene expression analysis by qPCR, the cDNA synthesis was performed using High-Capacity cDNA Reverse Transcription Kit with RNase Inhibitor (Cat. #4374966, Applied Biosystem).

For RNA-seq, RNA concentration was measured using a Qubit 2.0 Fluorometer (Life Technologies, Carlsbad, CA, USA), and RNA integrity was assessed with an Agilent TapeStation 4200 (Agilent Technologies, Palo Alto, CA, USA). RNA-seq libraries were prepared using the NEBNext Ultra II RNA Library Prep Kit for Illumina according to the manufacturer’s instructions (NEB, Ipswich, MA, USA). Briefly, polyadenylated mRNA was enriched using oligo(dT) beads and subsequently fragmented for 15 minutes at 94 °C. First- and second-strand cDNA synthesis was performed, followed by end repair and 3′ adenylation. Universal adapters were ligated to the cDNA fragments, and indexed libraries were generated by limited-cycle PCR amplification. Library quality was evaluated using the Agilent TapeStation, and library concentration was determined using a Qubit 2.0 Fluorometer (Invitrogen, Carlsbad, CA, USA) and qPCR (KAPA Biosystems, Wilmington, MA, USA).

Sequencing libraries were clustered on a flow cell and sequenced on an Illumina platform (HiSeq 4000 or equivalent) following the manufacturer’s protocols. Paired-end sequencing was performed with a read length of 2 × 150 bp. Image analysis and base calling were carried out using Illumina Control Software. Raw base call files were converted to FASTQ format and demultiplexed using bcl2fastq version 2.17, allowing one mismatch for index sequence identification.

### RNA-seq data analysis

RNA-seq data were processed using the Galaxy server [60]. Raw FASTQ files were first assessed for quality using FastQC, followed by adapter and low-quality base trimming when necessary. Transcript abundance was quantified directly from trimmed reads using Salmon v.0.8.2 [61] in quasi-mapping mode against the Akata-EBV (KC207813.1) and human (GRCh38) combined transcriptome. Differential expression analysis was then performed using edgeR 3.34.0+galaxy [62], including library size normalization, dispersion estimation, and statistical testing between experimental conditions. Genes with adjusted *p* values below 0.05 were defined as differentially expressed (**see Table S2**).

### *In situ* proximity ligation assay

Proximity ligation assay (PLA) was performed as previously described [58, 63]. Briefly, Akata (EBV+) cells were blocked with 3% BSA in PBS for 1 hour at room temperature, then incubated overnight at 4□°C with either PBS control or a mixture of rabbit anti-HNRNPA2B1 (Cat. #14813-1-AP, Proteintech) and mouse anti-KDM1A (Cat. #67037-1-Ig, Proteintech) antibodies (1:50 in 3% BSA). Probes were subsequently incubated at 37□°C for 1 hour, followed by ligation and signal amplification using the NaveniFlex Cell Red kit (Cat. #NC.MR.100, Navinci). Nuclei were counterstained with DAPI, and samples were imaged on a Nikon AXR confocal microscope.

### Quantification and statistical analysis

Statistical analyses were performed using a two-tailed Student *t*-test with Microsoft Excel software. A *p* value less than 0.05 was considered statistically significant. The values are presented as means and standard deviations for biological replicate experiments as specified in the figure legends.

## DATA AVAILABILITY

The RNA-seq data sets have been deposited in the NCBI Gene Expression Omnibus (GEO) under accession number GSE320084 (**pending to be released**). The proteomics data sets have been deposited in the ProteomeXchange Consortium via the PRIDE [64] partner repository with identifier PXD074669 (**pending to be released**).

## ACKNOWLEDGEMENTS

We thank S. Diane Hayward (Johns Hopkins University) for providing reagents, plasmids, and cell lines. We are grateful to Didier Trono (EPFL) for pMD2.G and psPAX2 plasmids (Addgene #12259 and #12260), Feng Zhang for lentiCRISPR V2 vector TFORF2771-KDM5A and TFORF2768-KDM5D (Addgene #52961, #143614 and #142159), Nicolas Fawzi for MBP-hnRNPA2 (Addgene #98662), Monika Golas for pIDS-LSD1 (Addgene #109157) and Stephen Tapscott for pZLCv2-3xFLAG-dCas9-HA-2xNLS vector (Addgene #106357). We also thank George Tsao (University of Hong Kong) for providing HK-1 (EBV+) cells. We appreciate Shou-Jiang Gao, Yuang Chang, Haitao Guo, and Kathy Shair for insightful discussions.

This work was supported by the National Institute of Allergy and Infectious Diseases (grants AI141410 and AI187186) awarded to R.L., a Research Scholar Grant from the American Cancer Society (134703-RSG-20-054-01-MPC), and additional support from the University of Pittsburgh Medical Center, Hillman Cancer Center Grant P30CA047904 from National Cancer Institute, Virginia Commonwealth University Philips Institute for Oral Health Research, and the VCU Presidential Quest for Distinction Award. The funders had no role in study design, data collection and analysis, decision to publish, or preparation of the manuscript.

Conceptualization: R.L. and F.G.S.;

Data curation: F.G.S. and R.L.;

Formal analysis: F.G.S. and R.L.;

Funding acquisition: R.L.;

Investigation: F.G.S., Y.L, and R.L.;

Methodology: F.G.S. and R.L.;

Project administration: R.L.;

Resources: R.L.;

Supervision: R.L.;

Validation: F.G.S. and R.L.;

Visualization: F.G.S. and R.L.;

Writing (original draft): F.G.S.;

Writing (review and editing): R.L. and F.G.S.

